# Transcriptomic signature of rapidly evolving immune genes in a highland fish

**DOI:** 10.1101/822866

**Authors:** Chao Tong, Miao Li, Kai Zhao

## Abstract

Recent genome-wide studies have begun to elucidate the genomic basis of hypoxia, long-term cold and high saline and alkaline adaptation in highland fish, and a number of key genes contributed to its highland adaptation were identified. An increasing number of studies indicated that immune genes of Tibetan endemic fish species underwent positive selection towards functional shift, while the insight into immune gene repertoire of Tibetan highland fishes from genome-wide studies has largely lagged behind. In this study, we performed one of the first comparative genomics study in particular focusing on the signatures of immune genes in a highland fish, *Gymnocypris przewalskii* based on immune-relevant tissue transcriptome assemblies. We identified seven putative rapidly evolving immune genes with elevated molecular evolutionary rate (dN/dS) relative to lowland fish species. Using tissue-transcriptome data, we found most of rapidly evolving immune genes were broadly expressed in head-kidney, spleen, gills and skin tissues, which significantly enriched for complement activation and inflammatory response processes. In addition, we found a set of complement activation related genes underwent accelerated evolution and showed consistently repressed expression patterns in response to parasite *Ichthyophthirius multifiliis infection*. Moreover, we detected a number of immune genes involved in adaptive immune system exhibited distinct signature of upregulated expression patterns after parasite infection. Taken together, this study provided putative transcriptomic signatures of rapidly evolving immune genes, and will gain the insight into Schizothoracine fish adaptation to high-altitude extreme aquatic environments including diversified pathogen challenge.

## 1. Introduction

Understanding how fish species adapt to the aquatic life at high altitude is an interest for both ichthyologists and evolutionary biologists [1]. As the largest and highest highland in the world with the average elevation above 4000m [2], the Tibetan Plateau imposes extremely harsh aquatic environments on endemic fish fauna [3], including hypoxia [4], chronic cold [5,6] and high salinity and alkalinity [7,8]. Most endemic Tibetan fish species had developed unique morphological, physiological or genetic features to tolerate such harsh living conditions [6,9]. The Schizothoracinae is the largest and most diverse taxon of ichthyofauna on the Tibetan Plateau, which are distributed throughout the Tibetan Plateau and its peripheral regions [3,10]. Past evidences revealed that Schizothoracine fishes had well adapted to above extremely inhospitable aquatic environments [3,11,12]. In addition, recent genome-wide studies have begun to elucidate the genomic basis of hypoxia, long-term cold and high saline and alkaline adaptation in Schizothoracine fish [5–7,13,14]. Besides, the aquaculture of several Schizothoracine fish species has been developing rapidly due to the high economic values [15], such as *Gymnocypris przewalskii*. However, continuous growth of the aquaculture industry is threatened by infectious disease, mainly known as the “white spot disease”, which caused by infection with a highly virulent ciliate parasite *Ichthyophthirius multifiliis* [15–17]. The severe outbreaks of parasite infection could result in even more than 65% mortality and caused great economic losses in farmed *G. przewalskii* [18]. Therefore, these evidences indicated that Schizothoracine fish might have not well established the immune system against diversified pathogens invasion. It may provide novel insight into the highland adaptation by focusing in particular on the mechanism of immune genes in Schizothoracine fishes.

Recent studies employing genome-wide approaches identified massive immune genes in several aquaculture fish species [19,20]. With the recent rapid advances in sequencing technologies for non-model organisms without reference genomes, an increasing number of transcriptome-wide studies mainly focused on the identification of immune genes in fish species [21]. While, the insight into immune gene repertoire of Schizothoracine fishes from genome-wide studies has largely lagged behind [22,23]. Notably, a growing body of studies provided the evidences of immune genes showing signatures of rapid evolution and positive selection in Schizothoracine fishes, including *Gymnocypris przewalskii* [16,23,24] and *Gymnocypris eckloni* [25–27]. Among Schizothoracine fishes, Tibetan naked carp (*Gymnocypris przewalskii*) is one of the best characterized Tibetan endemic fish species, and a number of key genes contributed to its highland adaptation were identified by previous studies [7,13,14,16,12,4,28]. Therefore, it is an interesting issue to investigate the evolutionary signature of immune genes that contributed to highland adaptation in Schizothoracinae using *G. przewalskii* as a case study in highland fish species.

In this study, we performed the comparative genomics analysis and identified the putative rapidly evolving immune genes in a highland fish, *G. przewalskii*. Specifically, we used recently available *G. przewalskii* transcriptome and other four teleost fish genomes for comparison. In addition, using tissue-transcriptomics, we also characterized the expression pattern of rapidly evolving immune genes in four immune-relevant tissues and response to parasite infection.

## 2. Materials and methods

### 2.1. Data collection and transcriptome assembly

We downloaded the available transcriptomic data of *G. przewalskii* from NCBI SRA database (https://www.ncbi.nlm.nih.gov/sra). Specifically, we used four unpublished tissue transcriptome sequencing data including gills, head-kidney, spleen and skin of *G. przewalskii*, plus a spleen transcriptome in response to *Ichthyophthirius multifiliis* infection [17]. Sequencing reads were checked for quality using package FastQC. Sequencing adapters and reads with a quality score < 20 were trimmed with Trimmomatic [29], resulting in clean reads. Transcriptome was *de novo* assembled with Trinity [30] with default parameters. After assembly, the redundant transcripts were removed using CD-HIT [31] with the threshold of 0.90, and only the longest transcript under each cluster was extracted as unigene (unique gene). Next, the open reading frame (ORF) of each unigene was predicted using MARKER [32] and TransDecoder (https://github.com/TransDecoder/TransDecoder).

### 2.2. Orthologous gene group identification

We included four well-annotated teleost fish genomes and downloaded from Ensembl database (release 96) (http://useast.ensembl.org/index.html) to build a local protein database, including zebrafish (*Danio rerio*), tilapia (*Oreochromis niloticus*), medaka (*Oryzias latipes*) and spotted gar (*Lepisosteus oculatus*). We then translated the nucleotide sequences of protein-coding genes from the transcriptome assemblies of *G. przewalskii* into amino acid sequences using an in-house-developed perl script, and pooled it into above exiting protein database. Next, we conducted a self-to-self BLAST search for all protein sequences using DIAMOND [33] with an E-value cutoff of 1e^−5^, and removed hits with identity < 30% and coverage < 30%. We identified the orthologous gene groups from the BLAST results using OrthoFinder [34] with default settings. We calculated, mapped and illustrated all the identified orthologous gene groups of each species by venn diagram. Finally, we identified the core orthologous gene groups shared by above five fish species.

Among the genes under above identified orthologous gene groups, we identified one to one, one-to-many, and many-to-many orthologs among five fish species. Notably, for each 1:1 ortholog pair, we only selected the longest transcript associated with the gene for each pair of species as putative single-copy ortholog. In addition, the one-to-many orthologs were treated as multiple copy orthologs, and many-to-many orthologs were labeled as other orthologs. Finally, the number of genes under every category in each fish species were calculated, respectively.

### 2.3. Phylogenetic inference

We performed the sequence alignment of each putative single-copy ortholog of five fish species using R package, MSA [35] with default parameters and trimmed gaps using trimAl [36] with parameter “-automated1”. Moreover, we performed the filtration of single-copy orthologs with stricter constraints, including length (minimum 200 aa), sequence alignment (maximum missing data 50% in CDS alignments). Next, we conducted a genome-scale coalescent-based dataset including thousands of single-copy genes from five fish species. Finally, we detected the best model for tree construction using ModelTest2 [37], and then we built the Maximum Likelihood phylogenetic tree using RAxML 8 [38].

Statistical supports for major nodes were estimated from 1000 bootstrap replicates. We reconstructed the species tree using ASTRAL 4.4.4 [39] for molecular evolution analysis.

### 2.4. Rapidly evolving gene identification

We sought to estimate the evolutionary rate of each single-copy ortholog in each fish species, and identified a set of genes with an elevated ratio of non-synonymous to synonymous substitutions (dN/dS) in *G. przewalskii* relative to other four fish species. At first, we derived nucleotide alignments from previously prepared protein alignments of each single-copy ortholog using PAL2NAL v14 [40]. We ran two branch models using CodeML package in PAML 4.7a [41] to identify rapidly evolving genes (REGs) in *G. przewalskii* lineage with corresponding nucleotide alignments, specifically with the null model assuming that all branches have been evolving at the same rate and the alternative model allowing the focal foreground branch (*G. przewalskii*) to evolve under a different evolutionary rate. Next, we used a likelihood ratio test (LRT) in R software, package MASS with df□=□1 to discriminate between the alternative model and the null model for each single-copy orthologs in the genesets. We only considered the genes as evolving with a significantly faster rate in the foreground branch if the adjusted *P* value□<□0.05 and higher dN/dS in the focal foreground branch than focal background branches (other four fish species). Finally, we annotated the rapidly evolving genes with gene ontology (GO) function category using R software, package topGO [42].

### 2.5. Gene expression analysis

We mapped all the clean sequencing reads to the transcriptome assemblies of *G. przewalskii* using RSEM [43] to obtain expected counts and transcripts per million (TPM). To avoid the contamination during dissection causes the low levels of expression in the neighboring structures, we removed genes with TPM < 1 in each dataset. We are primarily interested in the expression pattern of rapidly evolving immune genes (REIGs), and calculated the TPM value of each REIG in each tissue. At last, we annotated the differentially expressed REIGs by gene ontology using R software, package TopGO [42].

## 3. Results

### 3.1. Sequence analysis and orthologous gene groups

The *de novo* transcriptome assembly of *G. przewalskii* yielded 614,528 transcripts in total, with an N50 of 1,656 bp and an average length of 881 bp. After removing redundant isoforms and extraction of longest isoform among alternative transcripts, a total of 31,982 unigenes were obtained, with an N50 of 3,072 bp and a mean length of 1,985 bp. After prediction of protein-coding genes by MAKER and TransDecoder, we totally obtained 28,917 unigenes with full or partial length of gene coding regions (CDS) in *G. przewalskii* (supplementary table S1).

A total of 113,529 proteins from five fish species protein coding gene sets were binned into 30,218 orthologous gene groups. We identified a total of 10,334 core orthologous gene groups shared by eight fish species (Figure 1A). After strict one to one ortholog selection, we identified 5,573 putative single-copy orthologs (only one ortholog in each gene family) in each fish species among core shared orthologous gene groups (Figure 1B), making them suitable for phylogenetic inference.

**Figure 1.**
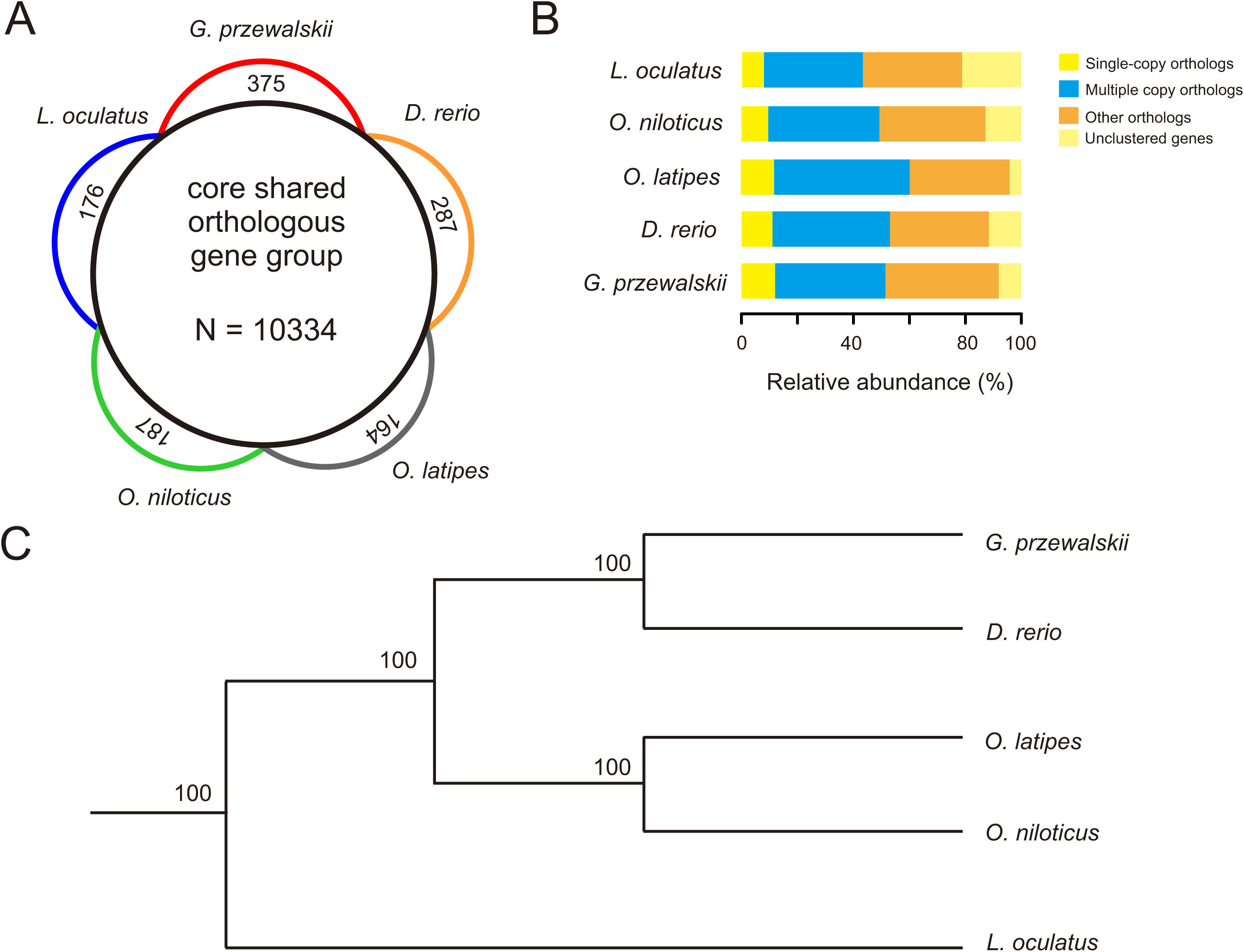
Genome content composition and genome-wide phylogeny. (A) Venn diagram showed the core shared orthologous gene groups (gene family) by five fish species. (B) Spine plot depicting the composition of different categories of gene families labeled by colors. (C) Genome-wide phylogeny of five fish species.

### 3.3. Phylogeny

After filtration for maximizing the information content of sequences and minimizing the impact of missing data, we eventually obtained 3,653 single-copy orthologs in each fish species. After protein sequence alignment, we constructed the maximum likelihood phylogenetic trees of five species based on coalesced single-copy orthologs using the GTRGAMMA model. The strongly supported (100%) phylogeny suggested that *G. przewalskii* had a closely related phylogenetic relationship with *Danio rerio*, and consistent with the tree on mitochondrial genes or nuclear DNA markers.

### 3.4. Rapidly evolving gene repertoire

Rapidly evolving gene is a set of genes underwent accelerated evolution with the signature of an increase rate of non-synonymous changes. We identified 468 putative rapidly evolving single-copy orthologs (REGs) in *G. przewalskii*. Among this set of genes, the most interesting finding was REGs included genes functioning in fish innate immune systems, such as Complement C2 (C2), Complement C5 (C5) and Complement factor D (CFD) that involved in complement activation process (Table 1). In addition, we found a number of REGs associated with fish adaptive immune response, such as Tumor necrosis factor alpha-induced protein 8 (TNFAIP8) and Tumor necrosis factor ligand superfamily member 10 (TNFSF10) involved in immune system process (Table 1). That is said, we identified a number of rapidly evolving immune genes in *G. przewalskii*. Besides immune genes, we also found a large number of genes involved energy metabolism functions, such as ATP5c1 and ATP5b associated with ATP binding and oxidative phosphorylation process (supplementary table S2).

**Table 1.**
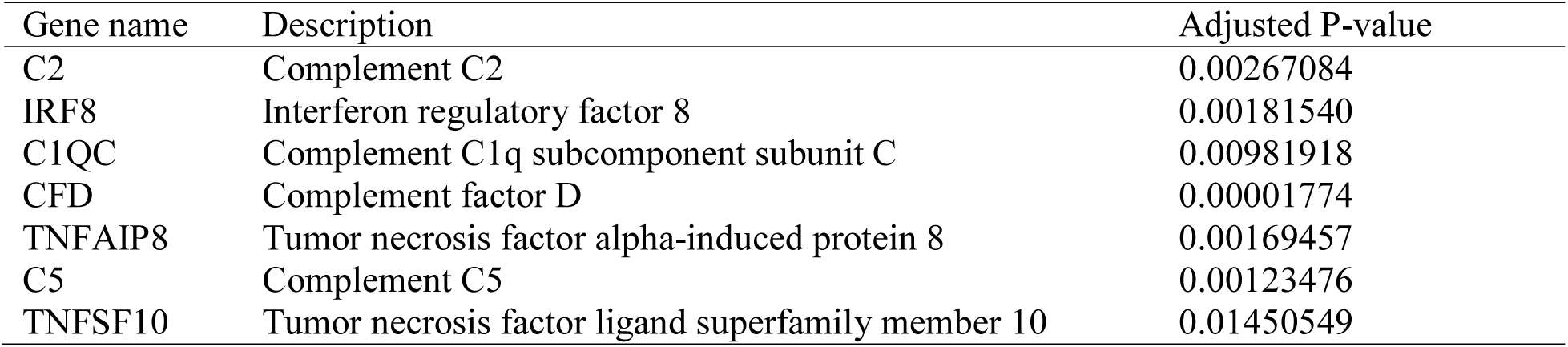
List of rapidly evolving immune genes in *G. przewalskii*.

| Gene name | Description | Adjusted P-value |
| --- | --- | --- |
| C2 | Complement C2 | 0.00267084 |
| IRF8 | Interferon regulatory factor 8 | 0.00181540 |
| C1QC | Complement C1q subcomponent subunit C | 0.00981918 |
| CFD | Complement factor D | 0.00001774 |
| TNFAIP8 | Tumor necrosis factor alpha-induced protein 8 | 0.00169457 |
| C5 | Complement C5 | 0.00123476 |
| TNFSF10 | Tumor necrosis factor ligand superfamily member 10 | 0.01450549 |

### 3.5. Expression patterns of rapidly evolving immune genes

We focused on the expression patterns of rapidly evolving immune genes (REIGs) among four immune-relevant tissues of *G. przewalskii*, including gills, head-kidney, spleen and skin. Most REIGs were broadly expressed in above four tissues except Complement 2 (Figure 2A, supplementary table S3), and mainly involved in innate immune response, immune system process and complement activation (Figure 2B, supplementary table S4). In addition, we screened the expression patterns of REIGs in response to parasite invasion using a set of spleen transcriptome data. We found that four complement genes were all repressed after *I. multifiliis* infection (Figure 2C, supplementary table S5), which mainly significantly enriched in complement activation and innate immune response process (Figure 2D, supplementary table S6). Moreover, we detected a set of REIGs involved in adaptive immune system including TNFAIP8 and TNFSF10 that were induced by parasite infection (Figure 2C, supplementary table S5), which significantly enriched in defense response to protozoan and type I interferon signaling pathway (Figure 2D, supplementary table S6).

**Figure 2.**
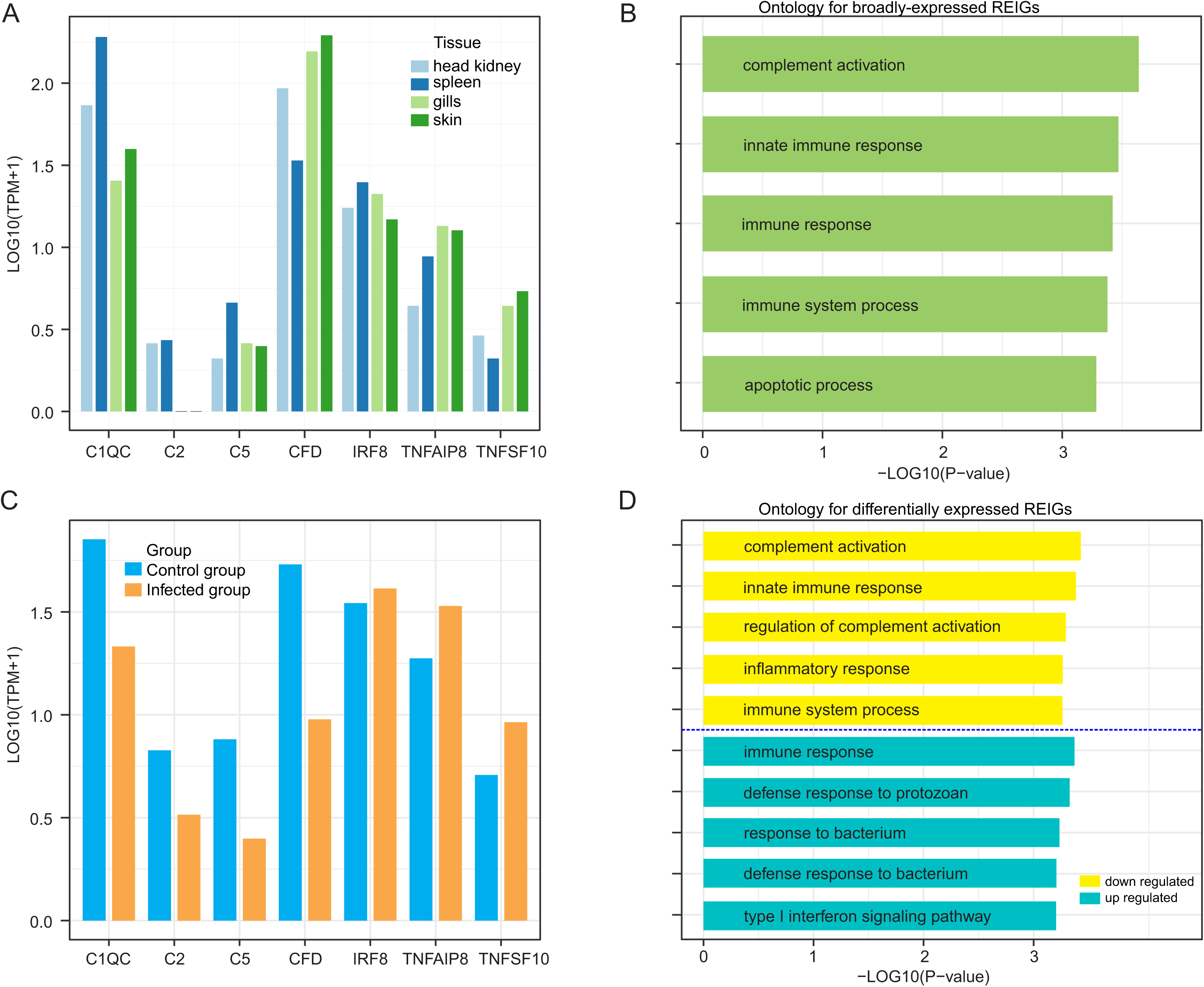
Expression patterns of rapidly evolving immune genes (REIGs) in four tissues, and spleen after parasite infection. (A) Bar plot depicting the expression level (Log_10_(TPM+1) value) of seven REIGs in head-kidney, spleen, gills and skin. (B) Bar plot depicting the top five gene ontology for broadly-expressed REIGs. (C) Bar plot depicting the expression level of seven REIGs in spleen between control and infected group. (D) Bar plot depicting the top five gene ontology for up- and down-regulated REIGs in *G. przewalskii*.

## 4. Discussion

Recent transcriptome-wide studies had begun to characterize a small scale of immune genes in Schizothoracine fishes [16,17,23,25], while our understanding of the genome-wide evolutionary signature of immune genes in highland fish remains limited. In the present study, for the first time, we used comparative genomics analysis to in particular identify rapidly evolving immune genes in Tibetan endemic fish, *G. przewalskii*. We found evidences of accelerated evolution acting on four complement genes involved innate immune system, two tumor necrosis factor genes associated with fish adaptive immune response process and one interferon regulatory factor gene. Using tissue transcriptome data, we found divergent expression patterns of rapidly evolving immune genes in head-kidney, spleen, gills and skin tissues of *G. przewalskii*. In addition, we found complement activation related immune genes underwent accelerated evolution exhibited distinct signature of downregulated expression in response to *Ichthyophthirius multifiliis* infection.

An increasing number of transcriptome targeted to identify abundant immune genes in fish by sampling immune-relevant tissues, such as head-kidney and spleen [21–23,25]. To maximize the information content of immune gene sequences, we collected two immune relevant tissues, head-kidney and spleen, plus two less relevant tissues (gills and skin) transcriptome data for assembly, which generated much more unigenes than our previous assembly versions of *G. przewalskii [7,23]*. In addition, compared with our previous comparative genomics study in *G. przewalskii* [7], we identified more immune genes showing signature of rapidly evolving relative to low land fish species. Unlike whole genome data, although transcriptome sequencing is an effective and accessible approach to initiate comparative genomic analyses on non-model organisms [30], it still can not cover all of the protein coding genes in one species. Moreover, most Schizothoracine fish species are polyploidy, the high complexity and large size of polyploidy fish genomes has constrained the development of genomic resources for Schizothoracinae [8]. The whole genome data of tetraploid *G. przewalskii* is long lacking, the limitation of transcriptome is obviously affecting the identification of immune gene repertoire. Therefore, it is not difficult to interpret that the increasing number of rapidly evolving immune gene were found in *G. przewalskii* due to including more immune-relevant tissue transcriptome data.

Accelerated evolution at molecular level may be reflected by an increased rate of non-synonymous changes within genes involved in adaptation [44]. Our present comparative genomics study identified a set of immune genes showing elevated evolutionary rate (dN/dS) in *G. przewalskii* relative to other low land fish species. We found four complement genes underwent accelerated evolution in *G. przewalskii*, such as C2, C1QC, CFD and C5. Complements are important components in fish innate immune system, and activation of complement can result in the generation of activated protein fragments that contributed to microbial killing, phagocytosis, inflammatory reactions, immune complex clearance, and antibody production [45]. Moreover, we found two rapidly evolving tumor necrosis factor (TNF) genes including TNFAIP8 and TNFSF10, this finding is identical to our previous findings [7,16,23,24]. TNF genes function in many immunological and inflammatory responses under either normal or pathological conditions [46]. Notably, recent study suggested that highland fish, salt-water-dwelling wild *G. przewalskii* was susceptible to infectious disease, which triggered high mortality in freshwater farming industry, such as parasite *Ichthyophthirius multifiliis* infection [15]. In addition, previous surveys on Tibetan lakes showed that hypersaline and alkaline Lake Qinghai had low diversity of pathogens [11,12], which indicated that wild *G. przewalskii* dwelt in a lighter pathogen load aquatic environment. Therefore, we speculated that immune genes of highland fish, *G. przewalskii* have experienced accelerated evolution and functional shifts to well adapted to this specific aquatic environment.

Fish head-kidney and spleen are two key tissues contributed to immune defense against pathogen invasion [47]. In present study, we found most of rapidly evolving immune genes (REIGs) were broadly expressed across all four tissues, indicating that immune genes experiencing accelerated evolution in *G. przewalskii* may have general functions, such as complement activation, inflammatory response and innate immune response. One interesting finding is that we found C2 gene was not expressed in *G. przewalskii* skin tissues, while this was not consistent with previous study in common carp (*Cyprinus carpio*) which indicated C2 was highly expressed in skin tissue [48]. Fish skin mucus is crucial to humoral immunity [49], the lack of complement gene expression in skin may affect the innate immune response of *G. przewalskii* to pathogen invasion. Moreover, it is interpretable that the accelerated evolution may drive complement genes of *G. przewalskii* towards functional shift during long-term adaptation to the low diversity of pathogens environment in Lake Qinghai. Another intriguing finding is that, all REIGs associated with complement activation function were repressed after *I. multifiliis* infection. Fish complement system is the first line of host defense, which promoted opsonization, lysis and local inflammatory response to pathogens infection [50]. Past evidences suggested that pathogens can avoid complement activation by mimicking host ligands, attracting complement regulators or secreting complement inhibitors [50–52]. In fact, explicit role of complement system was unclear, especially in response to *I. multifiliis* infection. Here, we speculated that *G. przewalskii* complement system may reduce the sensitivity to detect various pathogens during its adaptation to highland aquatic environment, this can result in this parasite *I. multifiliis* can evade or counteract its innate immune system to further trigger high mortality in freshwater farmed *G. przewalskii*. Since the complements were mainly synthesized in fish liver tissue, this hypothesis needs to be further confirmed by large-scale transcriptomics and functional assay in future.

## 5. Conclusion

We used comparative genomics based on the assemblies from multiple immune-relevant tissue transcriptomes to identify the signatures of immune genes a highland fish, *G. przewalskii*. These putative transcriptomic signatures included: 1) a number of genes involved in innate or adaptive immune system were found in *G. przewalskii* with elevated molecular evolutionary rate (dN/dS) relative to lowland fish species; 2) most rapidly evolving immune genes associated with complement activation and inflammatory response were broadly expressed in head-kidney, spleen, gills and skin of *G. przewalskii*; 3) A set of complement activation related genes underwent accelerated evolution and showed consistently repressed expression patterns in response to parasite *I. multifiliis* infection; 4) A set of genes involved in adaptive immune system exhibited distinct signature of upregulated expression after parasite infection. Taken together, our study will provide the transcriptomic signatures of rapidly evolving immune genes, and gain the insight into Schizothoracine fish adaptation to high-altitude extreme environments including diversified pathogen challenge.

## Supplementary Material

Supplementary table S1. Summary of protein-coding gene prediction.

Supplementary table S2. List of rapidly evolving genes in *G. przewalskii*.

Supplementary table S3. Expression levels of rapidly evolving immune genes in *G. przewalskii* head-kidney, spleen, gills and skin tissues.

Supplementary table S4. Ontology for broadly expressed rapidly evolving immune genes in *G. przewalskii*.

Supplementary table S5. Expression levels of rapidly evolving immune genes in spleen tissue *G. przewalskii* after parasite infection.

Supplementary table S6. Ontology for differentially expressed rapidly evolving immune genes in *G. przewalskii* spleen after parasite infection.

## Acknowledgments

This work was funded by the Field Research Fund of University of Pennsylvania Biology Department.

